# K-mer based classifiers extract functionally relevant features to support accurate Peroxiredoxin subgroup distinction

**DOI:** 10.1101/387787

**Authors:** Jiajie Xiao, William H. Turkett

**Affiliations:** Department of Computer Science, Wake Forest University, Winston Salem, NC 27109; Department of Physics, Wake Forest University, Winston Salem, NC 27109; Center for Molecular Signaling, Wake Forest University, Winston Salem, NC 27109

**Keywords:** Peroxiredoxins, Classification, Support vector machine, K-mer, Protein annotation

## Abstract

**Background:** The Peroxiredoxins (Prx) are a family of proteins that play a major role in antioxidant defense and peroxide-regulated signaling. Six distinct Prx subgroups have been defined based on analysis of structure and sequence regions in proximity to the Prx active site. Analysis of other sequence regions of these annotated proteins may improve the ability to distinguish subgroups and uncover additional representative sequence regions beyond the active site.

**Results:** The space of Prx subgroup classifiers is surveyed to highlight similarities and differences in the available approaches. Exploiting the recent growth in annotated Prx proteins, a whole sequence-based classifier is presented that employs support vector machines and a k-mer (k=3) sequence representation.

Distinguishing k-mers are extracted and located relative to published active site regions.

**Conclusions:** This work demonstrates that the 3-mer based classifier can attain high accuracy in subgroup annotation, at rates similar to the current state-of-the-art. Analysis of the classifier’s automatically derived models show that the classification decision is based on a combination of conserved features, including a significant number of residue regions that have not been previously suggested as informative by other classifiers but for which there is evidence of functional relevance.

## Background

The Peroxiredoxins (Prx) represent a family of enzymes found in a wide range of organisms that act as antioxidant defenses and play a role in managing cell signaling mediated by peroxide [1]. A highly conserved cysteine plays a primary role in imparting peroxide sensing functionality. Previous work [2,3] has provided evidence for six distinct subgroups of the Prx family - AhpE, Prx1-AhpC, Prx5, Prx6, PrxQ-BCP, and Tpx - and over 38,000 proteins have been annotated to the level of a Prx subgroup [4].

To discover proteins belonging to Prx subgroups, the state-of-the-art approach extracts sequence fragments containing residues within the active site region, a 10 angstrom region around the active site of Prxs for which structures are known. Subgroup specific active site sequence profiles can then be aligned against sequences in biological databases, with high scoring matches indicating likely membership of a protein into a subgroup. MISST [4], which implements a version of this search process that can iteratively expand and split clusters representing subgroups, resulted in the most recent 38,739 Prx subgroup-specific annotations.

These annotations approaches have, to date, limited sequence analysis to the active site regions. The idea of a k-mer representation, a list of counts of each length *k* sliding fragment along a sequence, has been widely adopted for fast approximations of a sequence identity [5,6,7,8], including applications to protein classification [9]. Subgroup-distinguishing k-mers represent small sequence fragments conserved between proteins within a subgroup but distinct across subgroups. It is shown in this manuscript that training a machine learning classifier on 3-mer-encoded subgroup-annotated proteins can allow for accurate Prx subgroup annotation. The distinguishing k-mers represent both previously known active site sequence fragments as well as additional sequence regions that are likely to be functionally relevant.

## Methods

### Data acquisition

The Structure-Function Linkage Database (SFLD) [10] provides a highly-curated protein database, organizing proteins by shared chemical function and providing a mapping between a given chemical function and associated active site features as represented in available protein sequences and structures. The SFLD, as of December 2017, annotates 7,345 proteins to the level of one of six Peroxiredoxin subgroups and annotates 12,239 (including the 7,345 annotated to the subgroup level) proteins as members of the Peroxiredoxin Superfamily.

The recent work of Harper *et al*. [4], using an iterative approach to search Genbank for proteins that have active site regions similar in sequence to those of known Prx structures, suggests a total of 38,739 Prx proteins annotated to a subgroup, of which 6,909 overlap with proteins annotated in the SFLD. The full data set of 38,739 proteins will be referred to as the *Harper* dataset, and the overlap set will be hereafter referred to as the *Harper-SFLD* dataset. The proteins in each subgroup of the Harper-SFLD data set were clustered at 95% sequence similarity using the CD-Hit algorithm [11, 12] to remove instances of proteins with high sequence similarity. The resulting set of 4,751 proteins will be referred to as the *0.95-Harper-SFLD* data set. The distributions over subgroups of the proteins in these data sets are shown in Table 1. There is imbalance among the subgroups, with up to an order of magnitude difference in the number of examples between subgroups.

**Table 1:** Counts of proteins in each Prx subgroup in each dataset

| **Data Set** | **AhpE** | **Prx1** | **Prx5** | **Prx6** | **PrxQ** | **Tpx** | **Total** |
| --- | --- | --- | --- | --- | --- | --- | --- |
| Harper | 1,489 | 9,660 | 5,434 | 5,212 | 12,014 | 4,930 | 38,739 |
| Harper-SFLD | 152 | 2,130 | 1,039 | 942 | 1,786 | 860 | 6,909 |
| 0.95-Harper-SFLD | 138 | 1,310 | 725 | 702 | 1,330 | 546 | 4,751 |

Each column represents the number of examples for a Prx subgroup available in the corresponding data set. The Harper-SFLD data set is the result of the intersection of the Harper data set with the subgroup-annotated Peroxiredoxins available in SFLD as of December 2017. The 0.95-Harper-SFLD data set encompasses the representative proteins after clustering the Harper-SFLD data set using the CD-Hit algorithm with a 95% similarity setting.

### Model construction

3-mers were used to encode protein sequences. With 20 amino acid residue options at each position of the 3-mer, this leads to 8,000 potential 3-mer features. Six one-versus-all classifiers were constructed, one per subgroup. All classifiers were built using linear-kernel support vector machines (SVM). Rather than other supervised learning methods, support vector machines (SVM) with linear kernels were chosen due to their effectiveness and efficiency in problems with high-dimensional features [13, 14]. The SVM technique optimally identifies the maximum-margin hyperplane that separates the positive and negative classes in feature space [15]. Given the fact that the features for the developed classifier SVM are k-mer occurrences, the linear-kernel utilized is also referred to as a *spectrum kernel* in the literature for binary classifications on biological sequences [8, 9].

The SVM-Light toolkit [16] was used for training and classification, with default values, automatically chosen by the SVM-Light implementation, used for training parameters. To classify a given protein sequence, the subgroup annotation associated with the maximum of the scores returned from the six classifiers was used. The Peroxiredoxin 3-mer SVM Classifier constructed will be hereafter referred to as Prx_3-merSVM.

## Results

### Classifier performance

Ten-fold cross validation was performed on the 0.95-Harper-SFLD data set, with 100% accuracy obtained in the cross validation experiment. To allow for a comparison with the work of Harper *et al*., a classifier built on all sequences from the 0.95-Harper-SFLD data set was then employed to classify the samples in the 38,739 protein Harper data set. Note that this large data set contains the 4,751 examples used for training. None of these 4,751 were classified incorrectly and these counts have been removed from the rest of the presented results. The confusion matrix comparing annotations generated by the Harper technique to those generated by the Prx_3-merSVM approach is shown in Table 2.

**Table 2:** Confusion matrix for classification on Harper data set

|  | **AhpE** | **Prx1** | **Prx5** | **Prx6** | **PrxQ** | **Tpx** |
| --- | --- | --- | --- | --- | --- | --- |
| **AhpE** | 1,348 | 0 | 1 | 0 | 2 | 0 |
| **Prx1** | 0 | 8,350 | 0 | 0 | 0 | 0 |
| **Prx5** | 0 | 0 | 4,709 | 0 | 0 | 0 |
| **Prx6** | 0 | 0 | 0 | 4,509 | 1 | 0 |
| **PrxQ** | 0 | 0 | 0 | 0 | 10,684 | 0 |
| **Tpx** | 0 | 0 | 0 | 0 | 0 | 4,384 |

The confusion matrix represents results from testing on the Harper data set, minus the 4,751 proteins in that set used for training. For a given protein, the row represents the known annotation as per Harper *et al.* and the column represents the annotation suggested by the Prx\_3-merSVM classifier. The counts represent how many proteins had each pair of annotations, with large values along the diagonal, representing matching annotations, being ideal.

### Distinguishing k-mers

Using the subgroup models constructed from the complete 0.95-Harper-SFLD data set, an exemplar set of distinguishing 3-mers (shown in Table 3) for each subgroup were extracted. These 3-mers were selected based on the ordered weights of the features from the linear kernel SVMs trained for each subgroup and permutation testing to determine the significance of observing such weights. Complete lists of 3-mers ordered by weight for each subgroup are included in the additional file *Additional file 1*. For all subgroups other than Prx1, the top ten high-weight 3-mers are included. For Prx1, only the top seven 3-mers surpassed the permutation testing threshold.

**Table 3:**
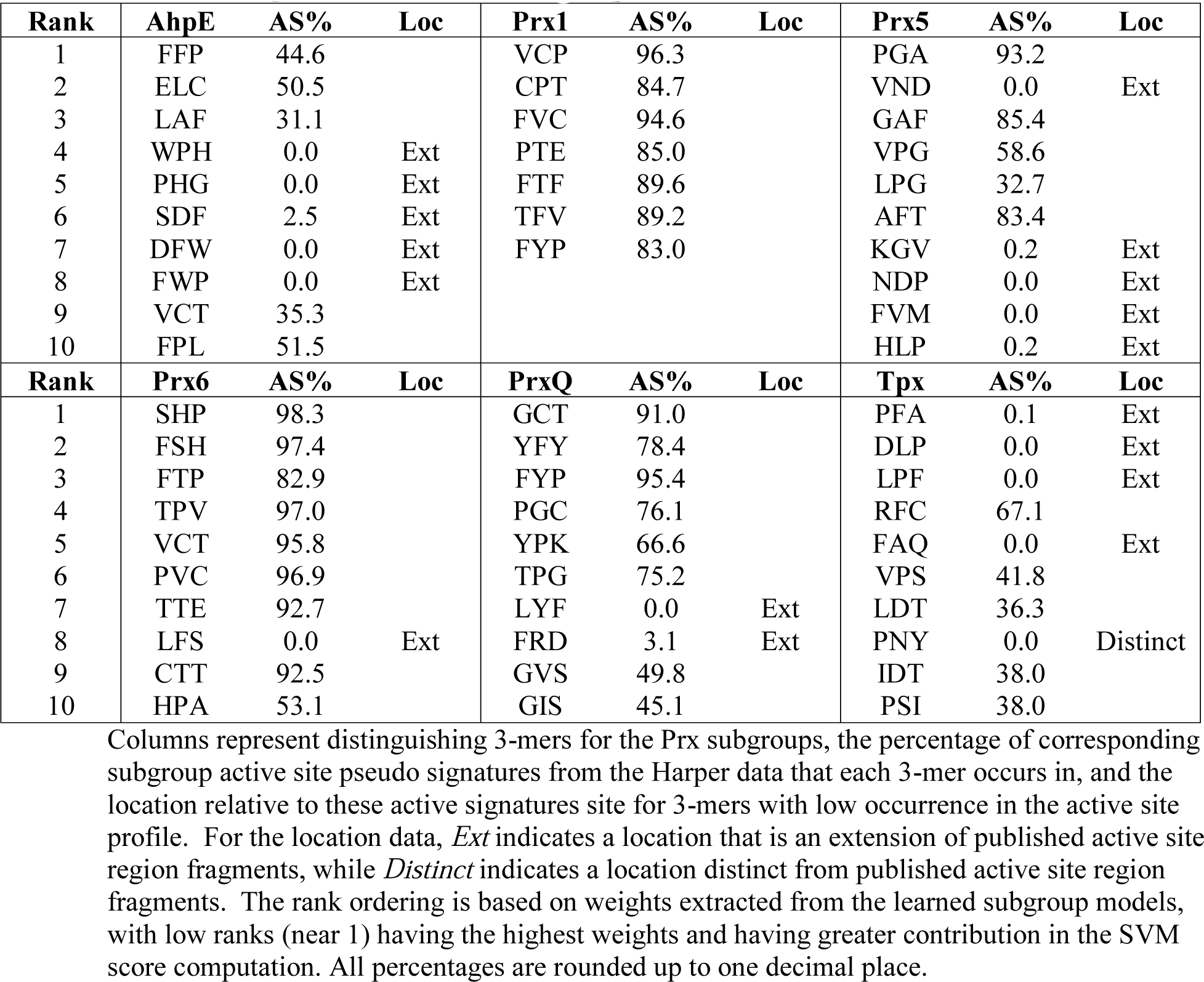
Discriminating 3-mers for each Prx subgroup

Permutation testing to determine a threshold SVM score at which to consider a SVM feature *highly discriminating* [17] was performed. For each Prx subgroup, the training data labels were randomly permuted and training was re-performed. This was repeated 2,000 times for each subgroup, allowing for an estimation of the distribution of SVM scores for each feature (3-mer) under a null hypothesis that there is no meaningful association between features and classes. The proportion of scores for a given feature greater than or equal to the observed score learned from the actual (non-permuted) training data was recorded as a P value. Given 2,000 permutations were performed, to ensure a conservative choice of 3-mers the highly discriminating features were constrained to only those whose score on the actual training data was greater than all scores on permuted data. All 3-mers in Table 3 are highly discriminating, and the permutation-testing based P value for all features is included in the supplementary material.

These 3-mers were searched for within the active site (pseudo-)signatures for the Harper data set proteins provided in the Supporting Information S2 file of [4]. Those signatures represent, for proteins identified to be members of each Prx subgroup, the sequence regions that best align with active site signatures for representatives of the subgroup, where an active site signature was defined by Harper *et al*. as the set of sequence fragments within 10 angstroms of the three selected active site residues. Repeated signatures for a subgroup were removed before searching for 3-mers. The search checked for whether a complete 3-mer was found as part of the signature sequence. The percentage of signatures for a subgroup in which each 3-mer is fully found is included next to 3-mers in the table. A significant proportion of active site region residues are represented by the distinguishing k-mers. This is particularly true for the Prx1 and Prx6 subgroups. These findings are reasonable given the high sequence conservation around the peroxidatic cysteine for these two subgroups as shown by Harper *et al*. It is also the case that a number of distinguishing k-mers are not contained within the published active site signatures. Through further sequence analysis, some resolve to extensions (nearby in sequence space) of the published sequence fragments, while others are new fragments in distinct parts of the sequence space. For 3-mers with low occurrence (less than 5%) in the pseudo-signature sequences, the location of the 3-mers is annotated in Table 3 as *Ext* if evidence suggests the 3-mer is an extension of the active site sequence fragments published by Harper *et al*. or as *Distinct* if the evidence suggests the 3-mer is in a distinct part of the sequence space. The determination of *Ext* or *Distinct* was made by extracting small regions of residues (8 residues in both directions) around the 3-mers of interest from the sequences containing the 3-mer, aligning the regions with ClustalOmega [18], generating a Weblogo [19], and visually inspecting the Weblogo against the Harper-specified regions. Information on 3-mers marked as *Ext* or *Distinct* that are not directly discussed in the manuscript is available in *Additional file 2*.

## Discussion

### Classification process comparison

With respect to the process of searching, the developed classifier has several advantages compared to other methods to classify Prx proteins at the subgroup level. Table 4 indicates the features of five different methods that can be used to classify Prx proteins to the subgroup level. These methods include HMM search against the SFLD database [10], use of the MISST algorithm [4] which builds on DASPs [20], BLAST search against the PREX database [3], search against the NCBI Conserved Domains database (CDD) [21], and the method described in this work named Prx_3-merSVM.

**Table 4:** Features of methods for annotating Prx proteins to the subgroup level

| Method | Web | Batch | Prx-specific | Hierarchical |
| --- | --- | --- | --- | --- |
| SFLD | Yes | No | No | Yes |
| MISST/DASP | No | Yes | Yes | No |
| PREX | Yes | No | Yes | No |
| CDD | Yes | Yes | No | Yes |
| Prx_3-merSVM | Yes | Yes | Yes | No |

Each row represents a method that can be employed to annotate Prx proteins at the subgroup level. Each column beyond the first represents a feature of a given method. Web represents whether or not a method is available via a web interface. Batch indicates whether more than one protein sequence can be processed at a time. Prx-specific indicates whether an annotation method is specific only to Prx subgroups. Hierarchical indicates whether the method is designed to return more generalized annotations in lieu of or in addition to a Prx subgroup annotation.

All methods except MISST/DASP2 have a web interface through which sequences can be uploaded to be analyzed. SFLD and PREX allow only one sequence to be analyzed at a time, reducing their utility for batch analyses, while MISST/DASP, CDD, and Prx_3-merSVM all support batch processing.

All of the approaches other than search against the PREX database are model-based, in that a model of subgroups is constructed and prediction is based on scoring against a model. These models are constructed via HMM learning in SFLD, construction of domain PSSMs with CDD, construction of active-site profiles with MISST/DASP, and construction of SVM models with Prx_3-merSVM. The PREX database process employs a BLAST search against its database of proteins. The sequence databases, models, and annotation techniques behind SFLD and CDD support the ability to provide annotations outside of the six Prx subgroups, including generalized annotations such as a Peroxiredoxin or Thioredoxin-fold annotation. The PREX database provide annotations to one of the Prx subgroups or indicates no annotation is appropriate, while Prx_3-merSVM assumes the protein is already known to be a Prx protein.

Prx_3-merSVM only currently provides as outputs the scores for each Prx subgroup for an input sequence. Other searching methods have hooks to additional information in their search output. Given PREX uses BLAST, it provides E-values for and alignments of the query sequence against high-scoring hits (hits with E-values less than 1E-40). The SFLD HMM search returns the level in the SFLD hierarchy for which an HMM match occurred, a corresponding E-value and score for the match, and the option to align against representative sequences in the matched group. CDD provides Prx subgroup annotations as specific conserved domain hits, with the output including RPS-BLAST E-values for the domain hits and the ability to see alignments against the sequences used to model a given domain. With the MISST/DASP approach, for a given subgroup, the relevant matching pseudo-signature, composed of the segments in the query sequence that best match the subgroup active site profile, and a DASP search score are output.

Several of the classification approaches also support means to directly query the data sets underlying their search mechanism. The PREX database can be searched using text via keyword, accession number, or genus and species. Proteins in the database associated with the search term are returned, including the associated Prx subgroup, the active site signature for the protein, and alignments against representative Prx proteins within the same subgroup and from other subgroups for comparison. The SFLD database can be searched by functional domain name, external identifiers, and by species, returning sequences, conserved residues, and the ability to compare the sequences against the rest of the SFLD database using BLAST and HMM approaches. In addition, sequence similarity networks for each Prx subgroup are available from SFLD, where edges between protein nodes are annotated with percent identity and logged E-value as computed by BLAST. Harper *et al*. provide in supplementary information active site region sequence signatures for all of the proteins that were annotated by their method.

### Classification performance comparison

The classification process employed by Harper *et al*. makes use of alignment against active site profiles (ASP) [20, 22], where an active site profile consists of multiple (usually 4 to 5) sequence fragments for which the residues are within 10 angstroms in structural space of known active site key residues. The Harper annotations [4] are considered the current gold standard for Prx annotations.

As a minimalist baseline to compare against the performance of the approaches described in Harper *et al*. and in this manuscript, six subgroup-specific canonical sequence motifs were also searched for in the proteins of the Harper data set. These six canonical active site motifs were chosen by reviewing the subgroup Weblogos provided by Harper *et al*., and then extracting the one sequence region in each that extended the general Prx active site motif PXXX(T/S)XXC. The motifs are shown in column 2 of Table 5. The data in that table represents for the Harper data set the percentage of proteins in each subgroup containing the simplified motif for that subgroup. These percentages align with the percentages reported by Harper [4], but are further broken down to show the percentage of sequences in each subgroup for which there are matches to more than one subgroup canonical motif. While searches for these canonical motifs alone would allow for accurate classification into some subgroups, there would be significant issues with respect to the AhpE and Prx5 subgroups. Due to the nature of those two motifs, most matches against the Prx5 motif also match the AhpE motif (only cases where Y is in the 4^th^ position of the Prx5 motif do not).

**Table 5:**
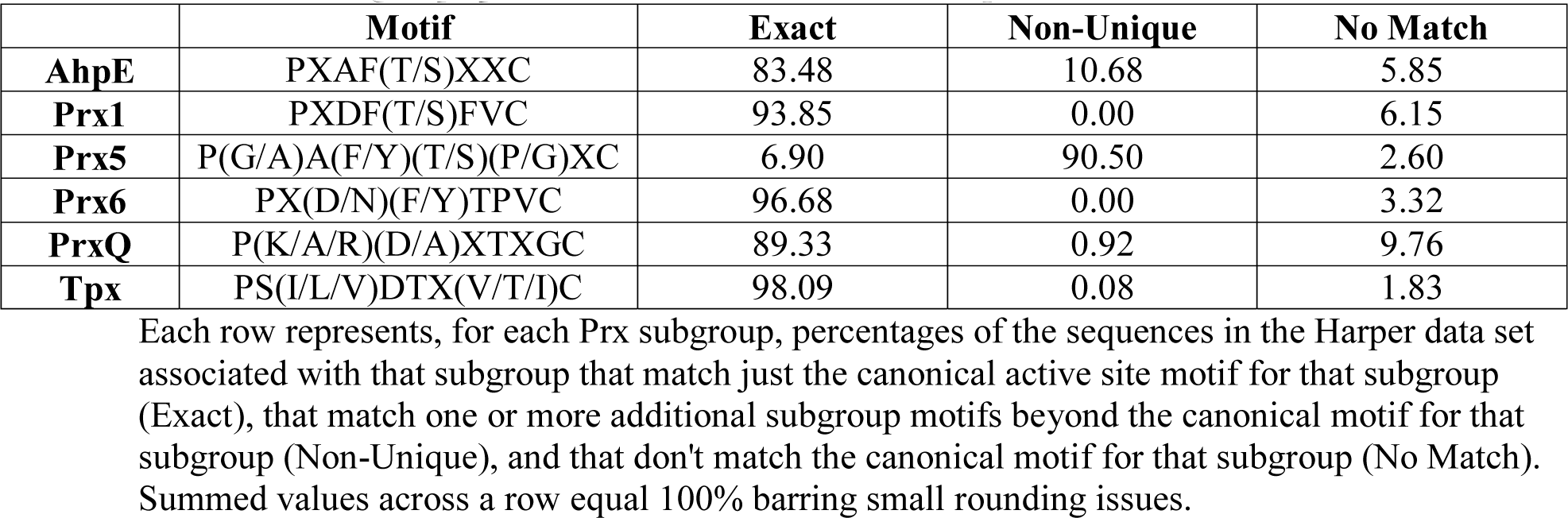
Matches of subgroup specific canonical motifs in the Harper data set

In Table 2, it was shown that there were four sequences that differed between the Harper and the 3-mer SVM annotations. These are described below by providing the Harper annotation, followed by the 3-mer SVM annotation. There was one instance of labeling an AhpE protein as Prx5, two instances of labeling an AhpE protein as PrxQ, and one instance of labeling a Prx6 protein as PrxQ. Additional annotations - from canonical motif search, the SFLD classifier, the PREX classifier, and the CDD classifier - for those four proteins are shown in Table 6.

**Table 6:**
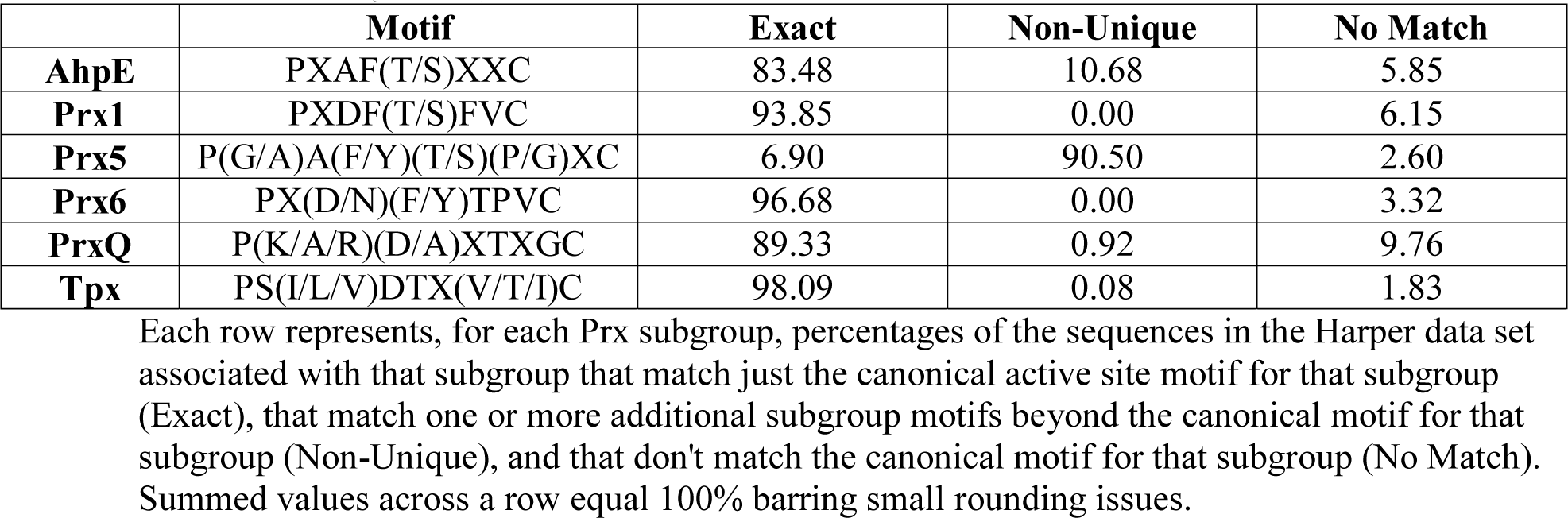
Annotations by multiple methods for proteins differing in annotation between Harper and Prx_3-merSVM

The developed classifier determines the subgroup to suggest for an input protein by selecting the annotation associated with the maximal score across the six subgroup classifier scores. For the four proteins with different annotations between the Prx_3-merSVM and Harper approaches, the Prx_3-merSVM scores from each subgroup classifier are shown in Table 7.

**Table 7:** Prx_3-merSVM classifier subgroup scores for proteins differing in annotation between Harper and Prx_3-merSVM

| Protein | AhpE | Prx1 | Prx5 | Prx6 | PrxQ | Tpx |
| --- | --- | --- | --- | --- | --- | --- |
| WP_055763280.1 | -0.606 | -1.054 | -1.137 | -1.002 | **0.498** | -1.121 |
| ELY63016.1 | -0.474 | -0.938 | **-0.430** | -0.897 | -0.669 | -0.915 |
| WP_051670221.1 | -0.353 | -0.949 | -0.774 | -1.220 | **-0.288** | -0.891 |
| KFD50172.1 | -0.883 | -0.896 | -0.438 | -0.714 | **-0.408** | -0.884 |

Each row represents a protein for which the Prx\_3-merSVM approach returned a label different from that provided by Harper [4]. The first column represents the Genbank id for a protein. The remaining columns provide the score returned from each subgroup specific model, with values rounded upwards. The maximum score for each protein is in bold font.

For the protein WP_055763280.1, considering the positive PrxQ score, the negative scores for the other subgroups, and the results from the PREX and CDD searches shown earlier, it is hypothesized that WP_055763280.1 actually belongs to the PrxQ subgroup. The other three proteins with differing annotations exhibit negative scores from all of the Prx_3-merSVM subgroup classifiers. Typically, the sign of the score returned from an SVM classifier can be used to indicate the class to which the given input belongs. A possible interpretation of all negative scores is that the proteins do not have characteristics of any of the subgroups. Reviewing the classifier outputs for the 38,739 protein Harper data set, negative scores were returned from all the subgroup classifiers for 63 of the proteins (including the three discussed above). Even with negative scores returned by all the subgroup classifiers, most of the Prx_3-merSVM annotations match the Harper annotation. The three differing annotations are from some of the lowest possible scores returned - these are shown as triangles in Figure 1. Similarly, the three proteins with differing annotations constitute three of the five proteins with the smallest difference between the highest scoring and next-highest scoring subgroup labels (for 38,739 proteins, that places them in the smallest 0.1%). For ELY63016.1 and WP_051670221.1 (the second and third proteins in Table 7), the Harper annotation is the second highest scoring Prx_3-merSVM annotation, but this does not hold for the fourth protein, KFD50172.1

**Figure 1:**
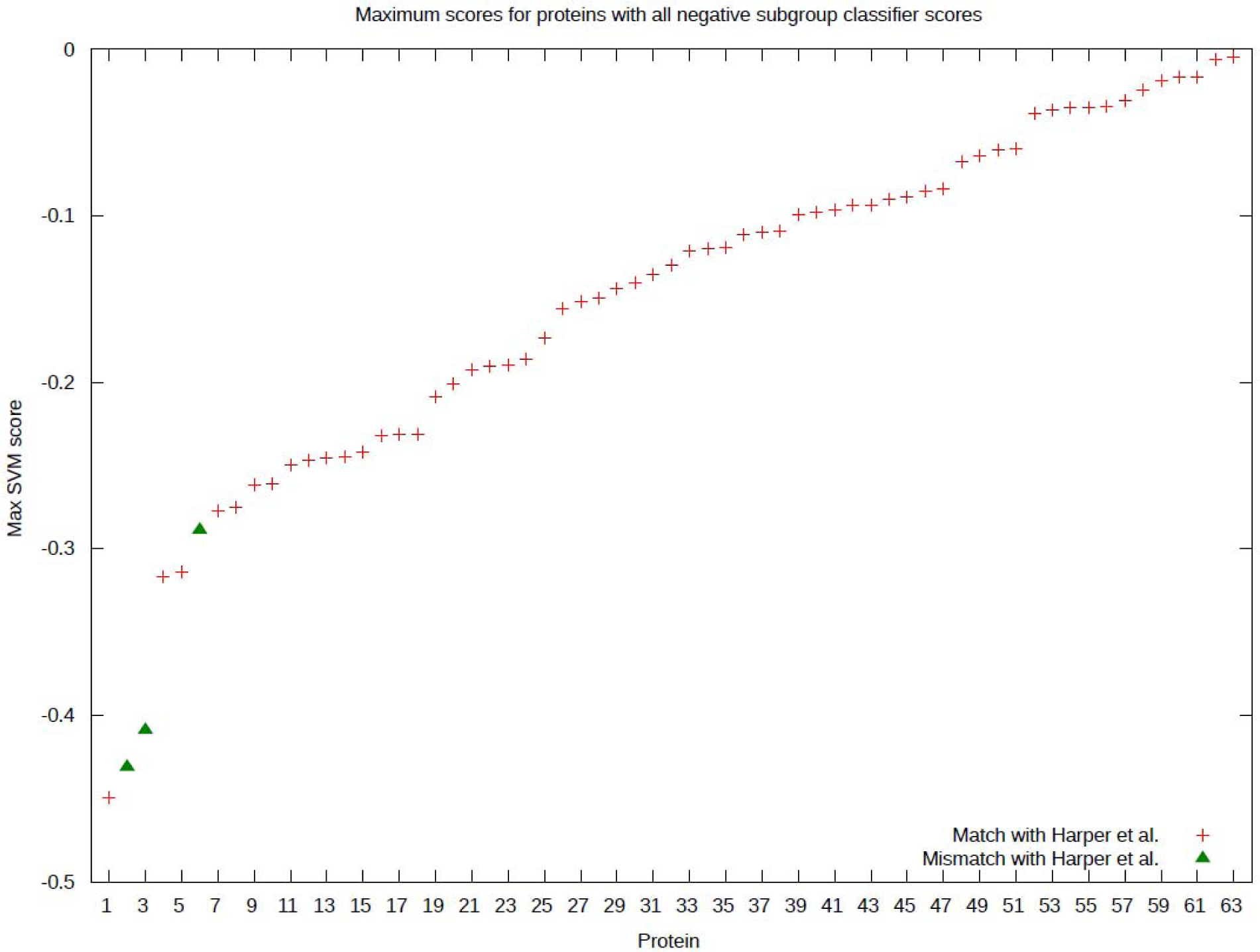
Maximum score for all proteins with negative scores from all Prx_3-merSVM subgroup
classifiers

Out of the 63 proteins with all negative Prx_3-merSVM scores, 53 are annotated as AhpE by Harper *et al*. 47 of those 53 are not in SFLD; the other 6 are in SFLD, but are not characterized to a subgroup. This is shown in Table 8. The AhpE subgroup has the least training data (an order of magnitude smaller than some of the other subgroups) and only has one structural representative. The signature conservation graph for AhpE in the work of Harper *et al*. is noisy relative to the other signature conservation graphs, highlighting increased variability in residues located structurally near the active site. Both of these help explain the lower-than-expected maximum scores for these proteins.

**Table 8:** Distribution of subgroup annotations for proteins receiving all negative Prx_3-merSVM scores

|  | AhpE | Prx1 | Prx5 | Prx6 | PrxQ | Tpx |
| --- | --- | --- | --- | --- | --- | --- |
| AhpE | 51 | 0 | 1 | 0 | 1 | 0 |
| Prx1 | 0 | 3 | 0 | 0 | 0 | 0 |
| Prx5 | 0 | 0 | 0 | 0 | 0 | 0 |
| Prx6 | 0 | 0 | 0 | 2 | 1 | 0 |
| PrxQ | 0 | 0 | 0 | 0 | 3 | 0 |
| Tpx | 0 | 0 | 0 | 0 | 0 | 1 |

The matrix represents the distribution of subgroup annotations for proteins from the Harper data set which received all negative scores from the Prx\_3-merSVM models. For a given protein, the row represents the known annotation (as per Harper *et al.*) and the column represents the annotation suggested by the developed classifier. The counts represent how many proteins had each pair of annotations.

### Analysis of distinguishing k-mers

Comparison of the distinguishing k-mers to the residues of Prx active sites suggests that a significant proportion of active site residues are represented by the distinguishing k-mers. In this work, as described previously, active site residues will be those annotated as being within an active site profile per Harper *et al*. Importantly, however, some distinguishing k-mers map in sequence space to functionally-relevant regions that are either extensions of the active site or are in distinct (non-active site) regions. Five exemplar sets of residues are presented below to highlight the type of information that can be extracted and made use of by the Prx_3-merSVM approach. Weblogo images of the +/-8 residue sequence regions surrounding a given 3-mer of interest are included. While some alignments end up being greater than nineteen residues in length (for example, due to a repeated use of a 3-mer in a sequence), all logos have been trimmed to only show the nineteen residue majority component of the alignment.

For the AhpE subgroup, the set of 3-mers DFW, FWP, WPH, and PHG commonly occur together. The information in Table 9 represents in how many proteins in the 0.95-Harper-SFLD data set and in the Harper data set each 3-mer occurs and how often they all occur in the same protein. These 3-mers commonly occur as a region of residues DFWPHG that occur as an extension of the active site profile region described as (F/A/Y)(P/D)(L/D)(L/F/V)(S/T/E/A) by Harper *et al*. The image in Figure 2 is a Weblogo representation of the region +/-8 residues centered on the 3-mer FWP extracted from the set of 1,055 AhpE sequences that all four 3-mers occur in. This set of residues is annotated as a turn in available protein structures for AhpE (1XVW, 4X0X) and has been suggested as playing an important role in the oligomerization interface [23].

**Figure 2:**
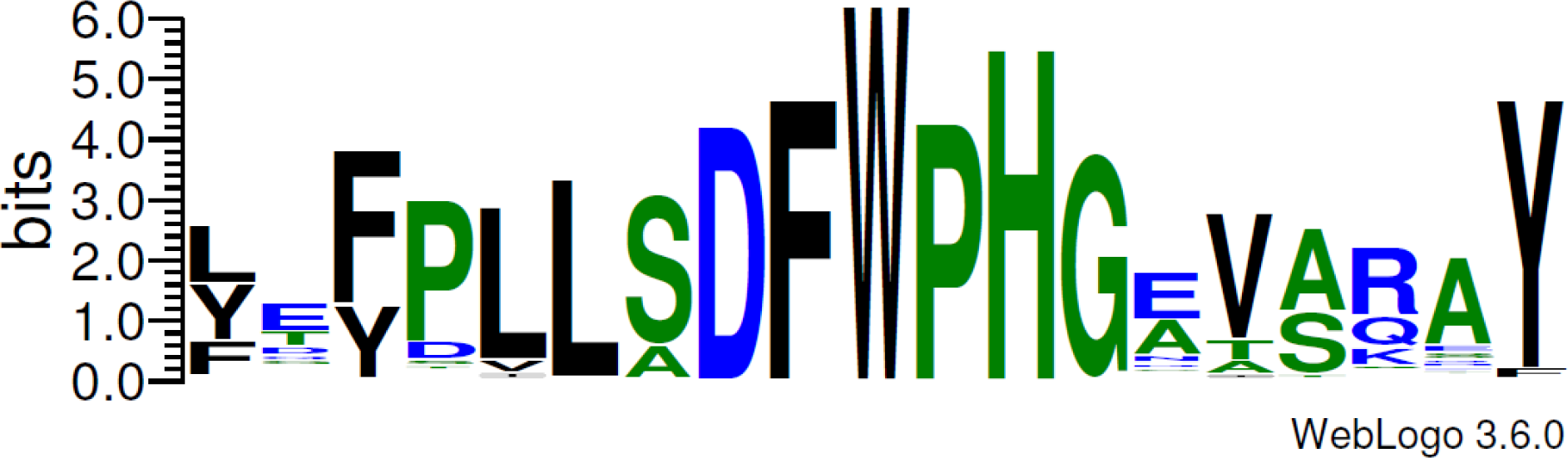
Weblogo of FWP-centered regions extracted from AhpE proteins

**Table 9:**
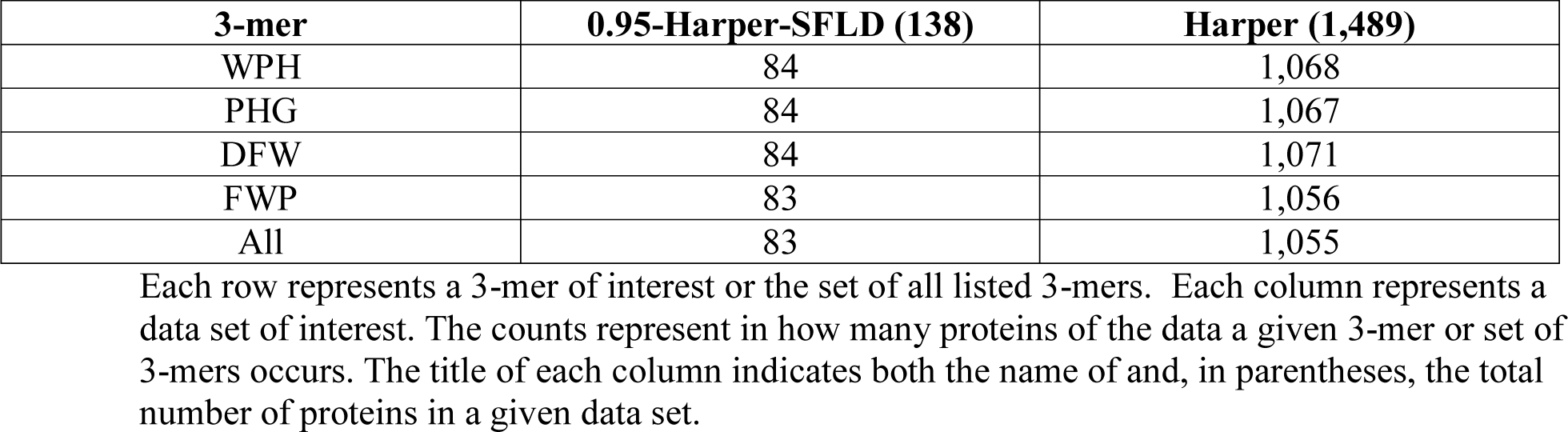
Counts of occurrence in AhpE proteins for four AhpE-distinguishing 3-mers

| **3-mer** | **0.95-Harper-SFLD (138)** | **Harper (1,489)** |
| --- | --- | --- |
| WPH | 84 | 1,068 |
| PHG | 84 | 1,067 |
| DFW | 84 | 1,071 |
| FWP | 83 | 1,056 |
| All | 83 | 1,055 |

Each row represents a 3-mer of interest or the set of all listed 3-mers. Each column represents a data set of interest. The counts represent in how many proteins of the data a given 3-mer or set of 3-mers occurs. The title of each column indicates both the name of and, in parentheses, the total number of proteins in a given data set.

For the Tpx subgroup, the set of 3-mers DLP, LPF, PFA, and FAQ commonly occur together. The information in Table 10 represents in how many proteins in the 0.95-Harper-SFLD data set and in the Harper data set each 3-mer occurs and how often they all occur in the same protein. These 3-mers commonly appear as an extension of the Tpx active site profile region described as A(Q/A/L/M)(K/A/S/G)R(F/W)C by Harper *et al*. The image in Figure 3 is a Weblogo representation of the region +/-8 residues centered on the 3-mer LPF extracted from the set of 3,570 Tpx sequences that all three 3-mers occur in. The set of residues corresponding with these 3-mers is annotated as a turn and the start of the alpha-helix containing the Tpx resolving cysteine in available protein structures for Tpx (1Y25, 3HVS). This region has been suggested as being highly conserved in sequence and playing roles as part of the dimer interface and as loop anchors [24].

**Figure 3:**
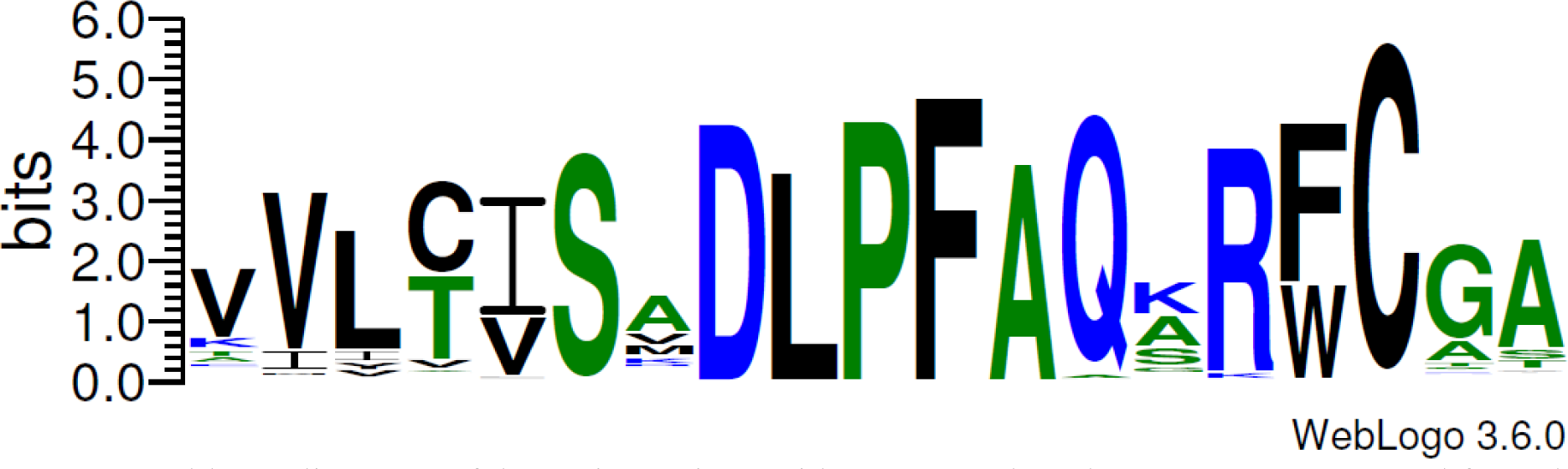
Maximum score for all proteins with negative scores from all Prx_3-merSVM subgroup classifiers

**Table 10:** Counts of occurrence in Tpx proteins for three Tpx-distinguishing 3-mers

| 3-mer | 0.95-Harper-SFLD (546) | Harper (4,930) |
| --- | --- | --- |
| DLP | 540 | 4,890 |
| LPF | 539 | 4,856 |
| PFA | 544 | 4,875 |
| FAQ | 356 | 3,602 |
| All | 351 | 3,570 |

Each row represents a 3-mer of interest or the set of all listed 3-mers. Each column represents a data set of interest. The counts represent in how many proteins of the data a given 3-mer or set of 3-mers occurs. The title of each column indicates both the name of and, in parentheses, the total number of proteins in a given data set.

The 3-mer PNY, also for the Tpx subgroup, is, out of the set of highly distinguishing 3-mers shown in Table 3, the one marked as *Distinct* from the active site sequence regions published in the work of Harper *et al*. In *Additional file 1*, PDY is the 11^th^ highly weighted 3-mer, also with a permutation testing P score of 0. PNY occurs in 233 of the 546 Tpx samples in the 0.95-Harper-SFLD data set and 2,275 of the 4,930 Tpx samples in the Harper data set, while PDY occurs in 253 of the 546 and 2,001 of the 4,930 respectively. The image in Figure 4 is a Weblogo representation of the region +/-8 residues centered on the 3-mer region P(N/D)Y as extracted from the set of 4,276 Tpx sequences containing either PNY or PDY. P(N/D)Y has been shown to be over 90% conserved across bacterial Tpx and to play a role in forming the structure of the cradle surrounding the α2 helix where the peroxidatic cysteine sits [24].

**Figure 4:**
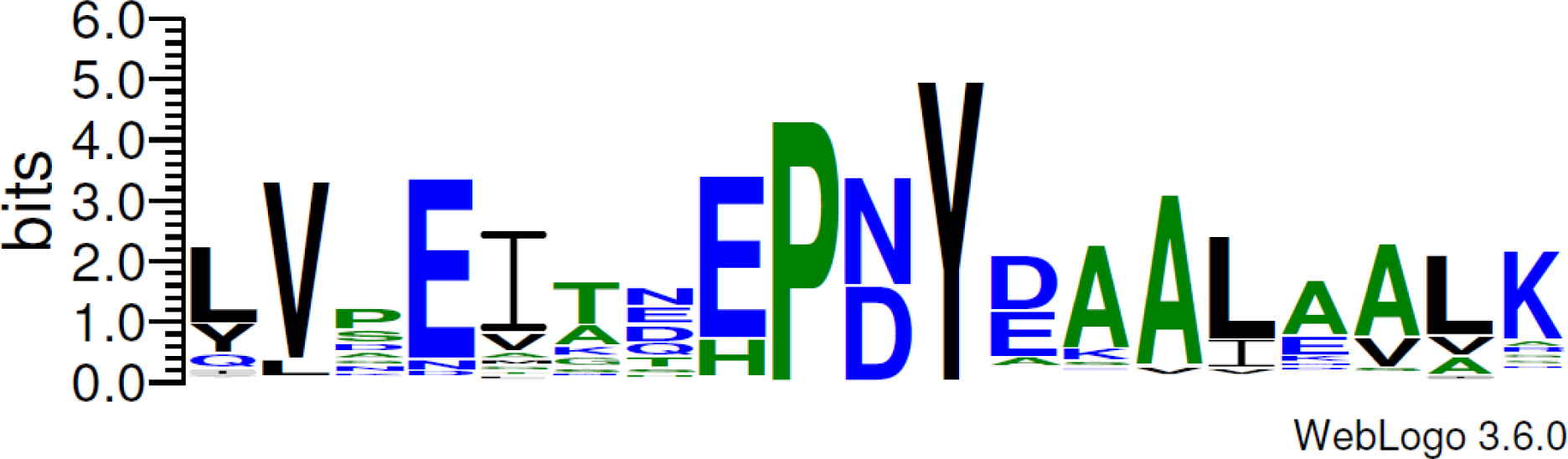
Weblogo of P(N/D)Y-centered regions extracted from Tpx proteins

For the Prx5 subgroup, the set of 3-mers VND and FVM occur together in a large percentage of the Prx5 protein sequences. The information in Table 11 represents in how many proteins in the 0.95-Harper-SFLD data set and in the Harper data set each 3-mer occurs and how often they all occur in the same protein. NDP is a less common extension of VND. VND and FVM are one-residue extensions to two different active sites regions described by Harper *et al*., the first region described as (C/V)(V/L/I/T/M)(S/A)VN and the other described by V(M/L/T)(N/G/K)(A/E/Q)W followed by several noisy positions. The high weights for VND and FVM suggests the D and F extensions are highly conserved as well. As shown in Figure 5 below, these two regions active site regions are themselves close in sequence space. They occur within or near the α3 helix (PDB 1NM3, 1HD2). Both the D and F residues are implicated as important in the dimerization interface [25] and the F residue (Phe79 in the around a benzoate ion ligand seen in the 1HD2 crystal structure [26].

**Figure 5:**
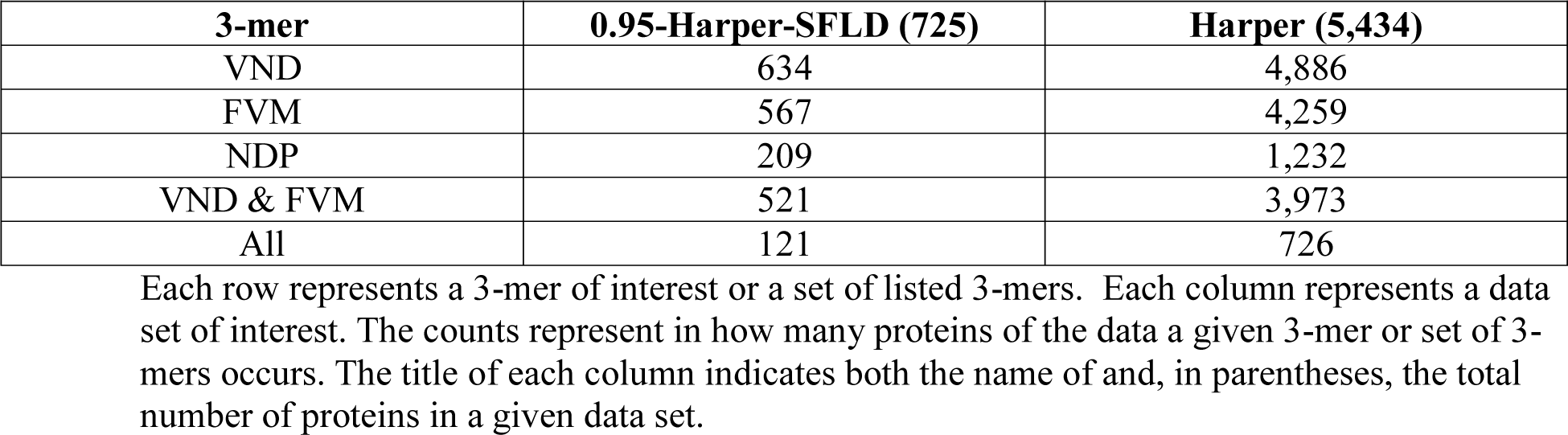
Weblogo of VND-centered regions extracted from Prx5 proteins

**Table 11:** Counts of occurrence in Prx5 proteins for three Prx5-distinguishing 3-mers

| 3-mer | 0.95-Harper-SFLD (725) | Harper (5,434) |
| --- | --- | --- |
| VND | 634 | 4,886 |
| FVM | 567 | 4,259 |
| NDP | 209 | 1,232 |
| VND & FVM | 521 | 3,973 |
| All | 121 | 726 |

Each row represents a 3-mer of interest or a set of listed 3-mers. Each column represents a data set of interest. The counts represent in how many proteins of the data a given 3-mer or set of 3-mers occurs. The title of each column indicates both the name of and, in parentheses, the total number of proteins in a given data set.

For the Prx1 subgroup, none of the highly distinguishing 3-mers (with permutation testing P score of 0) were in regions not seen by previous researchers. However, the 3-mer CPA, with P score 0.0035 (the most significant non-zero score; see *Additional file 1*), is an example of a region that is distinct from the active site sequence fragments published by Harper *et al*. CPA occurs in 1,065 of the 1,310 Prx1 samples in the 0.95-Harper-SFLD data set and 7,479 of the 9,660 Prx1 samples in the Harper data set. The image in Figure 6 is a Weblogo representation of the region +/-8 residues centered on the 3-mer CPA extracted from the set of 7,479 sequences the 3-mer occurs in. While distinct from the Harper active site sequence fragments, the C in this 3-mer represents the important resolving cysteine [27].

**Figure 6:**
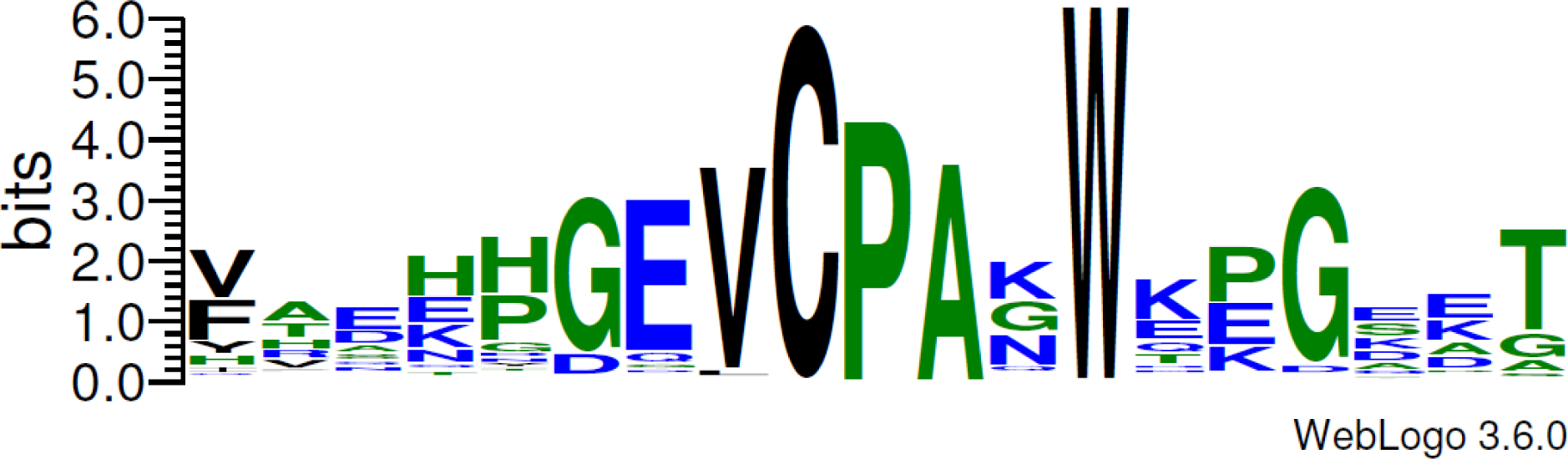
Weblogo of CPA-centered regions extracted from Prx1 proteins

### Limitations in analysis

This work demonstrates that the use of 3-mers supports high accuracy subgroup annotation of Prx sequences. The classifiers have been constructed under the assumption that a sequence to be annotated is already known to be a Prx sequence. To remove this constraint, the use of a hierarchical classification mechanism [28] could be developed to first annotate a protein as a Peroxiredoxin or not, and then to annotate to the subgroup level. A check for the presence of the Prx canonical active site motif PXXX(T/S)XXC could also play this role.

It is possible that a protein can receive negative scores from all six subgroup classifiers. A traditional approach to handling this scenario is to suggest that annotating the protein to one of the six subgroups is inappropriate when none of the scores is 0 or above. However, given the number of correct predictions made on the proteins in the Harper data set using the Prx_3-merSVM approach by using the annotation with the highest score, it may be suitable to adjust the threshold for when to suggest not providing an annotation to a score below 0.

While the discovered 3-mers highlight sequence regions that distinguish between Prx subgroups, the use of 3-mers is a fairly low resolution technique. A given 3-mer maps to a small portion of a given Prx sequence. The SVM classifier takes into account the presence of multiple 3-mers. The use of k-mers with larger k-values (4-mers, 5-mers) and the use of gapped k-mers [29], where wildcard (‘X’) positions are allowed in the k-mer, could potentially support accurate prediction with fewer and more interpretable features. The presence of several regions that could be captured by larger or gapped k-mers was highlighted in this work, including DFWPHG for the AhpE subgroup, DLPFAQ for the Tpx subgroup, and VNDXFVM for the Prx5 subgroup.

Additional members of the groups of highest weighted 3-mers, particularly those that are not part of the Harper published active site sequence regions, should be explored with respect to the role that the 3-mer residue regions play mechanistically. The use of other feature selection methods, such as recursive feature elimination (RFE) [30], to determine the subset of features to analyze beyond the exemplars provided is important, followed by analysis with respect to biochemical and biophysical features of the involved residues and the location of 3-mers in known protein structures.

Adding information beyond subgroup Prx scores to the output of the Prx_3-merSVM classifier would allow users to gain additional insight into their query proteins and the classification process.

### Conclusions

In this work, a new high-accuracy classifier that can annotate Prx proteins to the subgroup level has been developed. The classifier, which encodes sequences as 3-mers, is publicly available and supports batch analyses. Comparison to the state-of-the-art approach to Prx subgroup annotation shows only four differences in subgroup assignments in over 38,000 annotations. Examination of a subset of 3-mers that the developed classifier uses to distinguish between Prx subgroups reveals functionally relevant sequence fragments. These include sequence regions that extend or are distinct in sequence space from the active site sequence regions used in previous Prx subgroup analyses. Finding additional functionally-relevant regions has potential downstream uses in Prx inhibitor design.

## Declarations

### Ethics approval and consent to participate

Not applicable

### Consent for publication

Not applicable

### Availability of data and materials

The datasets generated and analyzed during the current study are available in the GitHub repository https://github.com/turketwh/Prx_3-merSVM_data, DOI: https://doi.org/10.5281/zenodo.1346271. Source code and an executable version of the software will be made available after a license for release has been developed by Wake Forest University. An Amazon Web Services hosted implementation of the classifier tool is publicly available at the address http://prxsubfamilyclassif-env.us-east-1.elasticbeanstalk.com/

### Authors’ contributions

WHT conceived the overall study and designed the experiments. JX and WHT performed the experiments, analyzed the data, and wrote the manuscript. All authors read and approved the final manuscript.

## Acknowledgements

The authors wish to acknowledge Dr. Leslie Poole for reviewing and providing helpful feedback and insights on initial drafts of the manuscript, as well as members of the Wake Forest University Center for Molecular Signaling for initial conceptual feedback. The authors acknowledge the Distributed Environment for Academic Computing (DEAC) at Wake Forest University for providing HPC resources that have contributed to the research results reported within this paper. URL: https://is.wfu.edu/deac WHT also wishes to acknowledge a Wake Forest University Reynolds Research Leave. JX acknowledges a fellowship from the Wake Forest University Center for Molecular Signaling.

## References

[1] Perkins A, Nelson KJ, Parsonage D, Poole LB, Karplus PA. Peroxiredoxins: guardians against oxidative stress and modulators of peroxide signaling. Trends Biochem Sci. 2015; 40(8):435–45.

[2] Nelson KJ, Knutson ST, Soito L, Klomsiri C, Poole LB, Fetrow JS. Analysis of the peroxiredoxin family: using active-site structure and sequence information for global classification and residue analysis. Proteins: Struct, Funct, Bioinf. 2011; 79(3):947–64.

[3] Soito L, Williamson C, Knutson ST, Fetrow JS, Poole LB, Nelson KJ. PREX: PeroxiRedoxin classification indEX, a database of subfamily assignments across the diverse peroxiredoxin family. Nucleic Acids Res. 2010; 39(D1):D332–7.

[4] Harper AF, Leuthaeuser JB, Babbitt PC, Morris JH, Ferrin TE, Poole LB, Fetrow JS. An atlas of peroxiredoxins created using an active site profile-based approach to functionally relevant clustering of proteins. PLoS Comput Biol. 2017; 13(2):e1005284.

[5] Altschul SF, Madden TL, Schäffer AA, Zhang J, Zhang Z, Miller W, Lipman DJ. Gapped BLAST and PSI-BLAST: a new generation of protein database search programs. Nucleic Acids Res. 1997; 25(17):3389–402.

[6] Jensen LJ, Gupta R, Blom N, Devos D, Tamames J, Kesmir C, Nielsen H, Staerfeldt HH, Rapacki K, Workman C, Andersen CA. Prediction of human protein function from post-translational modifications and localization features. J Mol Biol. 2002; 319(5):1257–65.

[7] Bhandare S, Goldberg DS, Dowell R. Discriminating between HuR and TTP binding sites using the k-spectrum kernel method. PLoS ONE. 2017; 12(3):e0174052.

[8] Fletez-Brant C, Lee D, McCallion AS, Beer MA. kmer-SVM: a web server for identifying predictive regulatory sequence features in genomic data sets. Nucleic Acids Res. 2013; 41(W1):W544–56.

[9] Leslie C, Eskin E, Noble WS. The spectrum kernel: A string kernel for SVM protein classification. Pac Symp Biocomput. 2002; 1:564–75.

[10] Akiva E, Brown S, Almonacid DE, Barber 2nd AE, Custer AF, Hicks MA, Huang CC, Lauck F, Mashiyama ST, Meng EC, Mischel D. The structure-function linkage database. Nucleic Acids Res. 2013; 42(D1):D521–30.

[11] Li W, Godzik A. Cd-hit: a fast program for clustering and comparing large sets of protein or nucleotide sequences. Bioinformatics. 2006; 22(13):1658–9.

[12] Fu L, Niu B, Zhu Z, Wu S, Li W. CD-HIT: accelerated for clustering the next-generation sequencing data. Bioinformatics. 2012; 28(23):3150–2.

[13] Pochet NL, Suykens JA. Support vector machines versus logistic regression: improving prospective performance in clinical decision-making. Ultrasound Obst Gyn. 2006; 27(6):607–8.

[14] Shao Y, Lunetta RS. Comparison of support vector machine, neural network, and CART algorithms for the land-cover classification using limited training data points. ISPRS J Photogramm Remote Sens. 2012; 70:78–87.

[15] Cortes C, Vapnik V. Support-vector networks. Mach Learn. 1995; 20(3):273–97.

[16] Joachims T. Making large-scale support vector machine learning practical. In: Schölkopf B, Burges CJC, Smola AJ, editors. Advances in Kernel Methods. Cambridge: MIT Press: 1999. p169–84.

[17] Mourao-Miranda J, Bokde AL, Born C, Hampel H, Stetter M. Classifying brain states and determining the discriminating activation patterns: support vector machine on functional MRI data. Neuroimage. 2005; 28(4):980–95.

[18] Sievers F, Wilm A, Dineen D, Gibson TJ, Karplus K, Li W, Lopez R, McWilliam H, Remmert M, Söding J, Thompson JD. Fast, scalable generation of high-quality protein multiple sequence alignments using Clustal Omega. Mol Syst Biol. 2011; 7(1):539.

[19] Crooks GE, Hon G, Chandonia JM, Brenner SE. WebLogo: a sequence logo generator. Genome Res. 2004; 14(6):1188–90.

[20] Leuthaeuser JB, Morris JH, Harper AF, Ferrin TE, Babbitt PC, Fetrow JS. DASP3: identification of protein sequences belonging to functionally relevant groups. BMC Bioinformatics. 2016; 17(1):458.

[21] Marchler-Bauer A, Bryant SH. CD-Search: protein domain annotations on the fly. Nucleic Acids Res. 2004; 32(W1):W327–31.

[22] Cammer SA, Hoffman BT, Speir JA, Canady MA, Nelson MR, Knutson S, Gallina M, Baxter SM, Fetrow JS. Structure-based active site profiles for genome analysis and functional family subclassification. J Mol Biol. 2003; 334(3):387–401.

[23] Li S, Peterson NA, Kim MY, Kim CY, Hung LW, Yu M, Lekin T, Segelke BW, Lott JS, Baker EN. Crystal Structure of AhpE from Mycobacterium tuberculosis, a 1-Cys peroxiredoxin. J Mol Biol. 2005; 346(4):1035–46.

[24] Hall A, Sankaran B, Poole LB, Karplus PA. Structural changes common to catalysis in the Tpx peroxiredoxin subfamily. J Mol Biol. 2009; 393(4):867–81.

[25] Kim SJ, Woo JR, Hwang YS, Jeong DG, Shin DH, Kim K, Ryu SE. The tetrameric structure of Haemophilus influenza hybrid Prx5 reveals interactions between electron donor and acceptor proteins. J Biol Chem. 2003; 278(12):10790–8.

[26] Declercq JP, Evrard C, Clippe A, Vander Stricht D, Bernard A, Knoops B. Crystal structure of human peroxiredoxin 5, a novel type of mammalian peroxiredoxin at 1.5 Å resolution. J Mol Biol. 2001; 311(4):751–9.

[27] Parsonage D, Nelson KJ, Ferrer-Sueta G, Alley S, Karplus PA, Furdui CM, Poole LB. Dissecting peroxiredoxin catalysis: separating binding, peroxidation, and resolution for a bacterial AhpC. Biochemistry. 2015; 54(7):1567–75.

[28] Silla CN, Freitas AA. A survey of hierarchical classification across different application domains. Data Min Knowl Disc. 2011; 22(1-2):31–72.

[29] Ghandi M, Lee D, Mohammad-Noori M, Beer MA. Enhanced regulatory sequence prediction using gapped k-mer features. PLoS Comput Biol. 2014; 10(7):e1003711.

[30] Guyon I, Weston J, Barnhill S, Vapnik V. Gene selection for cancer classification using support vector machines. Mach Learn. 2002; 46(1-3):389–422.

